# Hyaluronic acid turnover controls the severity of Cerebral Cavernous Malformations in bioengineered human micro-vessels

**DOI:** 10.1101/2023.03.27.534302

**Authors:** Teodor E. Yordanov, Marco A. E. Martinez, Tyron Esposito, Juliann B. Tefft, Larisa I. Labzin, Samantha J. Stehbens, Alan E. Rowan, Benjamin M. Hogan, Christopher S. Chen, Jan Lauko, Anne K. Lagendijk

## Abstract

Cerebral Cavernous Malformations (CCMs) are vascular lesions that predominantly form in blood vessels of the central nervous system (CNS) upon loss of the CCM multimeric protein complex. The endothelial cells (ECs) within CCM lesions are characterised by overactive MEKK3 kinase and KLF2/4 transcription factor signalling, leading to pathological changes such as increased EC spreading and reduced junctional integrity. Concomitant to aberrant EC signalling, non-autonomous signals from the extracellular matrix (ECM) have also been implicated in CCM lesion growth and these factors might explain why CCM lesions mainly develop in the CNS. Here, we adapted a three dimensional (3D) microfluidic system to examine CCM1 deficient human micro-vessels in distinctive ECMs. We validate that EC pathological hallmarks are maintained in this 3D model. We further show that key genes responsible for homeostasis of Hyaluronic Acid (HA), a major ECM component of the CNS, are dysregulated in CCM. Supplementing the ECM in our model with forms of HA that are predicted to be reduced, inhibits CCM cellular phenotypes, independent of KLF2/4. This study thereby provides a proof-of-principle that ECM embedded 3D microfluidic models are ideally suited to identify how changes in ECM structure and signalling impact vascular malformations.

## INTRODUCTION

Cerebral cavernous malformations (CCMs) are raspberry shaped vascular lesions that manifest primarily in postcapillary venules of the central nervous system (CNS). Lesions can develop sporadically or be a consequence of familial mutations. Both sporadic and familial cases are associated with loss of function (LOF) mutations in vascular endothelial cells (ECs) for one of three genes; *KRIT1* (*CCM1*), *OSM* (*CCM2*) or *PDCD10* (*CCM3*)^1,2^. CCM patients can develop a variety of symptoms depending on the number, size, and location of lesions. Symptoms include headaches, neurological deficits such as seizures and epilepsy, and haemorrhagic stroke^3–5^.

CCM proteins can assemble as a multimeric complex^6,7^ that regulates the activity of an endothelial specific Mitogen-Activated Protein Kinase (MAPK)-MEKK3 pathway^8–12^. This ectopic MEKK/ERK signalling in CCM lesions leads to excessive KLF2 and 4 transcription factor activity, which has been proven to be a key driver of the disease^13–16^. Once this pathway has been initiated, CCM lesion growth occurs by clonal expansion and incorporation of neighbouring wild-type ECs^17,18^. Wildtype incorporation is suggested to occur via non-cell autonomous factors. Indeed loss of CCM alters the ultrastructure of the ECM microenvironment^19–21^, and excessive degradation of the ECM proteoglycan Versican by Adamts5^16^ has been shown to directly contribute to CCM lesion growth^22^. CCM1 can also dampen β1-integrin activation by stabilising the integrin inhibitor ICAP1, which is essential for appropriate fibronectin remodelling^23,24^. Despite the appreciation that non-cell autonomous factors play an important role in CCM disease progression, it remains challenging to specifically modify and study such ECM components *in vivo*.

To facilitate analysis of ECM structure and signalling in CCM pathogenesis, we here adapted a three-dimensional (3D) micro-fluidic system^25,26^ to culture human CCM1 deficient micro-vessels. These vessels are grown under flow and in tuneable ECM hydrogels. We applied this model to grow CCM vessels in brain-mimetic matrices, enriched with Hyaluronic Acid (HA), the major structural component of the brain ECM^27,28^. HA molecules can vary greatly in size, ranging from very short HA oligomers (oHA) up to long polymers, referred to as high molecular weight (HMW) HA, of several MDa^29–31^. Notably, HA synthesising and degrading genes are amongst the most dysregulated genes in CCM lesions^32–34^. We validated that such transcriptional changes exist that would result in a reduced presence of low molecular weight (LMW) HA and excessive turnover of HMW HA in the ECM surrounding CCM lesions. Since distinct HA molecules are appreciated to activate differential cellular signalling^29–31^, altered HA composition of the ECM would likely impact CCM pathology. We used our CCM micro-vessel model to test this and identified that LMW HA inhibits pathological EC enlargement, whereas HMW HA improved EC junction integrity. Composite ECM containing both HMW and LMW HA created the most favourable environment with CCM phenotypes markedly reduced, suggesting that reduced presence of these HA forms in CCM disease might contribute to the formation of unstable and leaky lesions.

## RESULTS AND DISCUSSION

### Establishing a human CCM micro-vessels model to study ECM interactions

For this study we aimed to interrogate how changes in the ECM might alter the severity of EC phenotypes within CCM vasculature. First, we generated a CCM1/KRIT1 loss of function (LOF) model in human umbilical cord endothelial cells (HUVECs) using CRISPR/Cas9, further referred to as CCM1 LOF (Figure 1A). Growing these CCM1 LOF cells in a 2D setting revealed increased KLF4 expression in the EC nuclei (Figure 1B-C), validating CCM pathological signalling is activated^13,15^. CCM1 LOF ECs also displayed thinner cell-cell junctions with significantly reduced expression of the adherens junction proteins Vascular endothelial (VE)-cadherin and β-catenin (Figure 1D-E). By utilising VE-cadherin expression at the junctions, we further determined that CCM1 LOF ECs were larger and more elongated than their wild-type (WT) counterparts (Figure 1D, F and G). In agreement with previous studies^15,35,36^, CCM1 LOF ECs contained prominent (F)-actin rich stress fibres (Figure 1D-E) with elevated levels of phosphorylated Myosin Light Chain (pMLC S19 and pMLC T18/S19) (Supplementary Figure 1), showing these cells are under increased acto-myosin tension.

**Figure 1:**
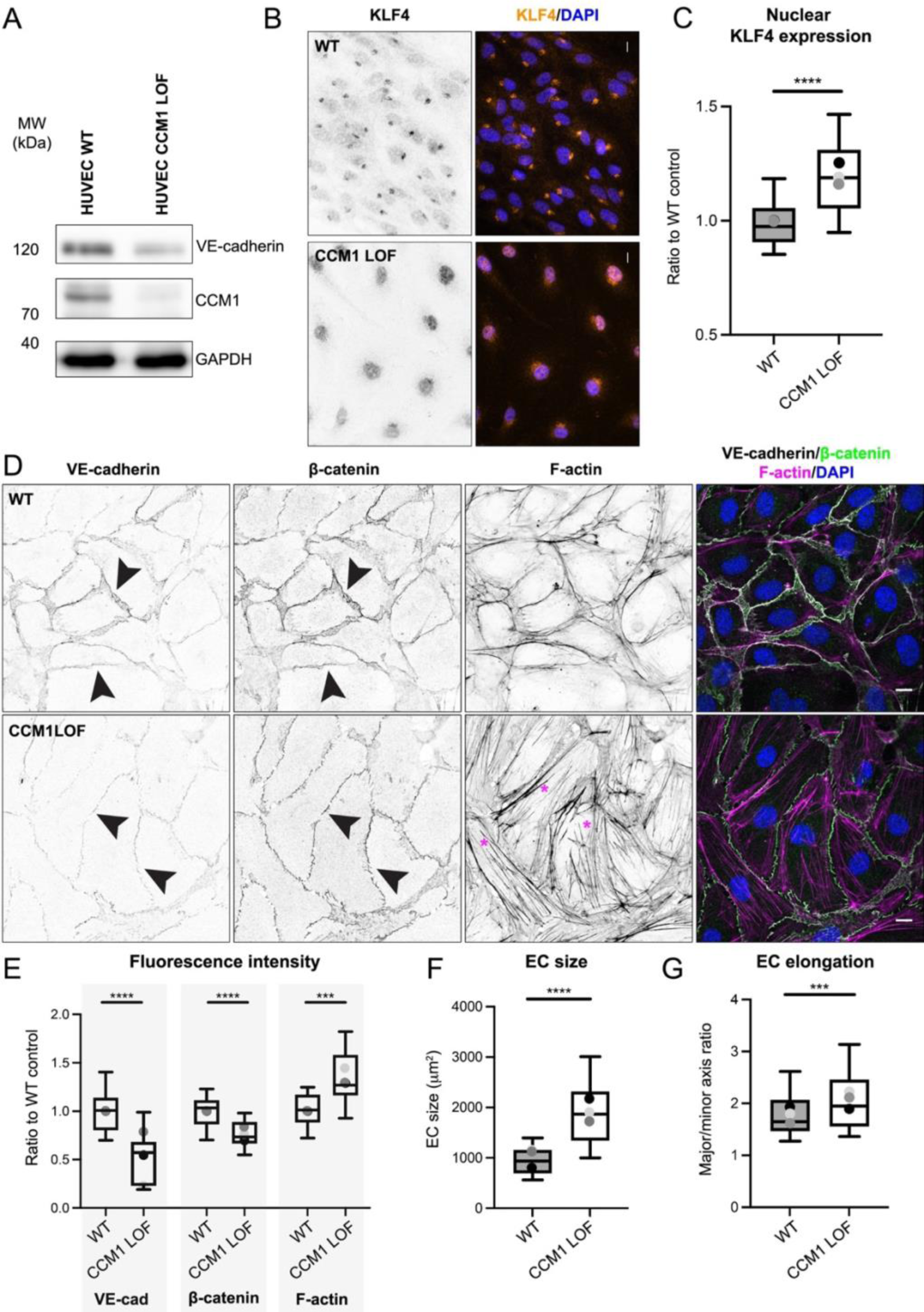
Generation and characterisation of CCM1 LOF HUVECs. (A) Western blot of WT and CCM1 LOF HUVECs showing loss of CCM1 (KRIT-1) protein expression. (B) Immunofluorescence of WT and CCM1 LOF ECs, stained with KLF4 and DAPI. Scale bar: 10 µm. Left panel: KLF only (grey). Right panel: KLF4 (orange) and DAPI (blue). (C) Box and whisker plot of KLF4 fluorescence intensity in the EC nuclei. Mean value of each independent replicate is represented as a dot with matching colours between WT and CCM1 LOF. n=3 replicates; n=150 WT and n=138 CCM1 LOF ECs, Mann-Whitney test ****p<0.0001. (D) Immunofluorescence of WT and CCM1 LOF ECs, stained for VE-cadherin, β-catenin, Phalloidin (F-actin) and DAPI. Arrowheads show adherens junctions, marked by VE-cadherin and β-catenin and asterisks (magenta) indicate F-actin stress fibres in CCM1 LOF ECs. Scale bar: 10µm. (E) Box and whisker plot indicating fluorescence intensity of VE-cadherin, β-catenin and Phalloidin (F-actin). Mean value of each replicate is represented as a dot with matching colours between WT and CCM1 LOF. n=3 replicates; n=18 WT and n=18 CCM1 LOF ECs, Student’s t-test ****p<0.0001 and ***p<0.001. (F) Quantification of the cell size of WT and CCM1 LOF ECs, based on VE-cadherin staining. Box and whisker plot with mean value of each replicate represented as a dot with matching colours between WT and CCM1 LOF ECs. n=3 replicates; n=150 WT and n=138 CCM1 LOF ECs, Mann-Whitney test ****p<0.0001. (G) Quantification of cell elongation (major axis over minor axis) of WT and CCM1 LOF ECs, utilising VE-cadherin to demarcate the cell boundaries. Box and whisker plot with mean value of each replicate represented as a dot with matching colours between WT and CCM1 LOF ECs. n=3 replicates; n=150 WT and n=138 CCM1 LOF ECs, Mann-Whitney test ****p<0.0001.

Next, we adapted a bio-engineered 3D model that was established by Polacheck and colleagues^25,26^ to grow CCM LOF micro-vessels. In this setup, the ECs are seeded on the luminal side of a 3D tube, surrounded by a physiologically relevant ECM, and placed under constant laminar flow pressure, thereby modelling the geometrical and environmental cues found *in vivo*. This setup is particularly suited to advance studies into the role of ECM in CCM pathogenesis.

We first seeded our validated CCM1 LOF ECs into a 3D Collagen-I ECM (2.5 mg/ml), the standardised ECM hydrogel used in this model^25,26^. Following the attachment of the ECs, the vessels were placed on a rocking platform to establish gravity-driven oscillatory flow^25,26^. We validated that CCM cellular characteristics were maintained in this 3D configuration (Movies 1 and 2), including decreased VE-cadherin coverage (Figure 2A-B), increased EC size and enhanced EC elongation (Figure 2C-D). We could identify these cellular changes as early as 24 hours post EC seeding, and these were maintained for the duration of our experiments (up to 5 days; Figure 2B-D). Together these experiments have validated that CCM1 LOF micro-vessels are a suitable model to study CCM biology. For reasons not well understood, CCM lesions form predominantly in the CNS. The ECM at these sites is known to be much softer compared to the ECM in many other tissues^37,38^. We therefore utilised our 3D models to determine whether an extremely soft environment would enhance CCM severity. We reduced the Collagen-I concentration in the 3D hydrogel from 2.5 mg/ml to 1.25 mg/ml. This resulted in a five-fold reduction of the ECM stiffness, from ∼400 Pa to 80 Pa, whilst maintaining the collagen structure and pore size (Supplementary Figure 3A-C). Quantifications of EC size (Figure 3A, B), junctional VE-cadherin expression (Figure 3A, C) and nuclear intensity of KLF4 (Figure 3A, D), uncovered a minor reduction in VE-cadherin whilst other CCM pathological phenotypes were unchanged. Together we have shown that major CCM cellular characteristics are maintained in a 3D geometry, whilst under flow, and in contact with a collagen-based ECM. Growing CCM vessels in a highly compliant ECM does not significantly alter phenotypic outcome, suggesting that low tissue stiffness in the CNS does not induce CCM formation.

**Figure 2.**
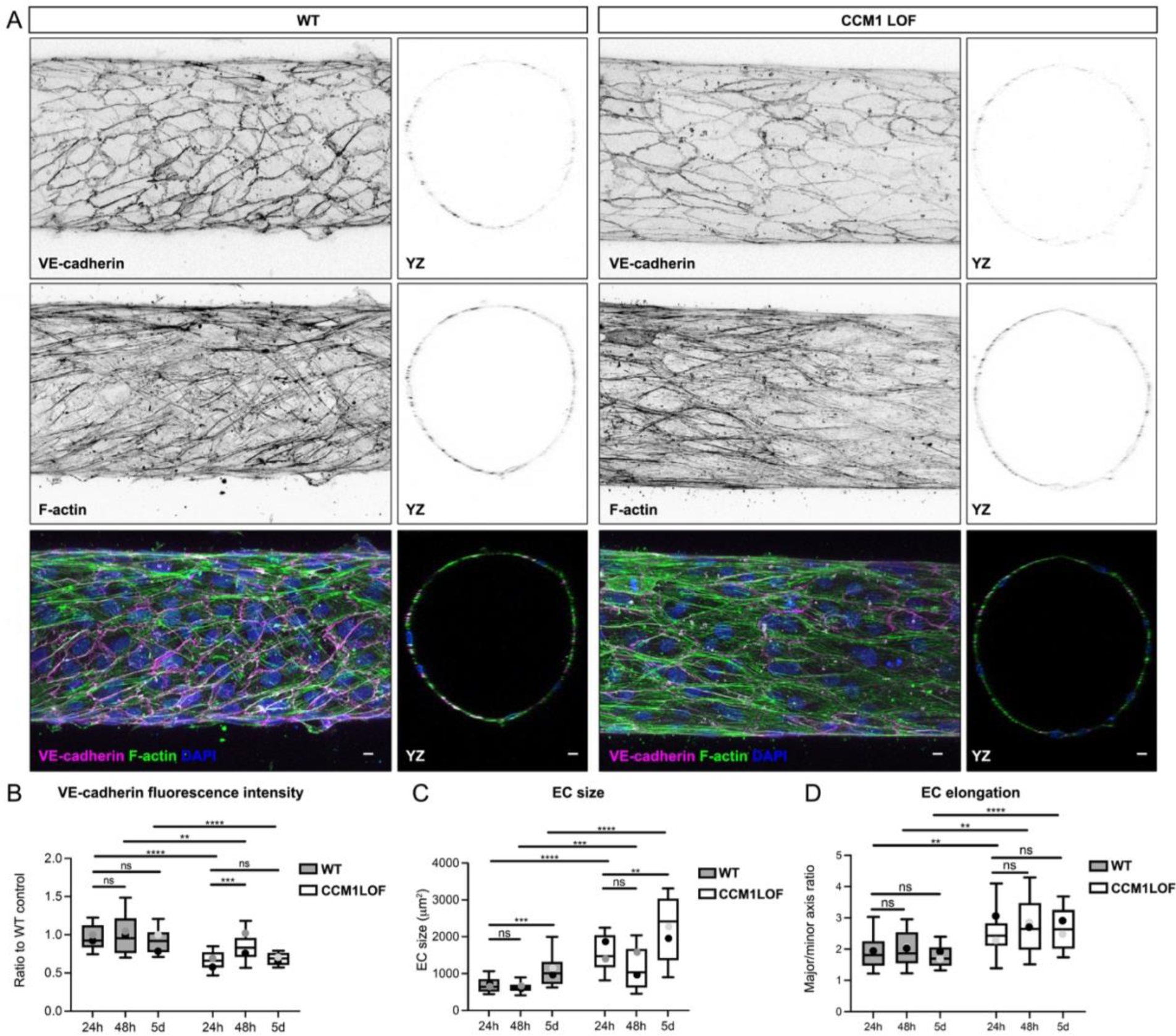
Establishing a human model of CCM deficient 3D vasculature. (A) Immunofluorescence of WT and CCM1 LOF EC cells, seeded in 3D microfluidic devices, grown for 48h post seeding, and stained for VE-cadherin (magenta), Phalloidin (F-actin) (green) and DAPI (blue). Images are presented as maximum projections in XY and YZ axis. Scale bar: 10µm. (B) Quantification of the VE-cadherin fluorescence intensity in ECs of WT and CCM1 LOF micro-vessels. Box and whisker plot with mean value of each replicate represented as a dot with matching colours between WT and CCM1 LOF ECs. n=3 replicates; 24h: n=39 WT and n=28 CCM1 LOF ECs, 48h: n=33 WT and n=27 CCM1 LOF ECs, 5d: n=24 WT and n=19 CCM1 LOF ECs, Student’s t-test ****p<0.0001, ***p<0.001, **p<0.005, ns=no significant difference. (C) Quantification of cell size of ECs in WT and CCM1 LOF micro-vessels. Box and whisker plot with mean value of each replicate represented as a dot with matching colours between WT and CCM1 LOF ECs. n=3 replicates; 24h: n=28 WT and n=22 CCM1 LOF ECs, 48h: n=33 WT and n=27 CCM1 LOF ECs, 5d: n=24 WT and n=17 CCM1 LOF ECs, Mann Whitney test ****p<0.0001, ***p<0.001, **p<0.005, ns=no significant difference. (D) Quantification of cell elongation (major over minor axis) of ECs in WT and CCM1 LOF micro-vessels. Box and whisker plot with mean value of each replicate represented as a dot with matching colours between WT and CCM1 LOF ECs. n=3 replicates; 24h: n=28 WT and n=22 CCM1 LOF ECs, 48h: n=33 WT and n=27 CCM1 LOF ECs, 5d: n=24 WT and n=17 CCM1 LOF ECs, Student’s test ****p<0.0001, **p<0.005, ns=no significant difference.

**Figure 3.**
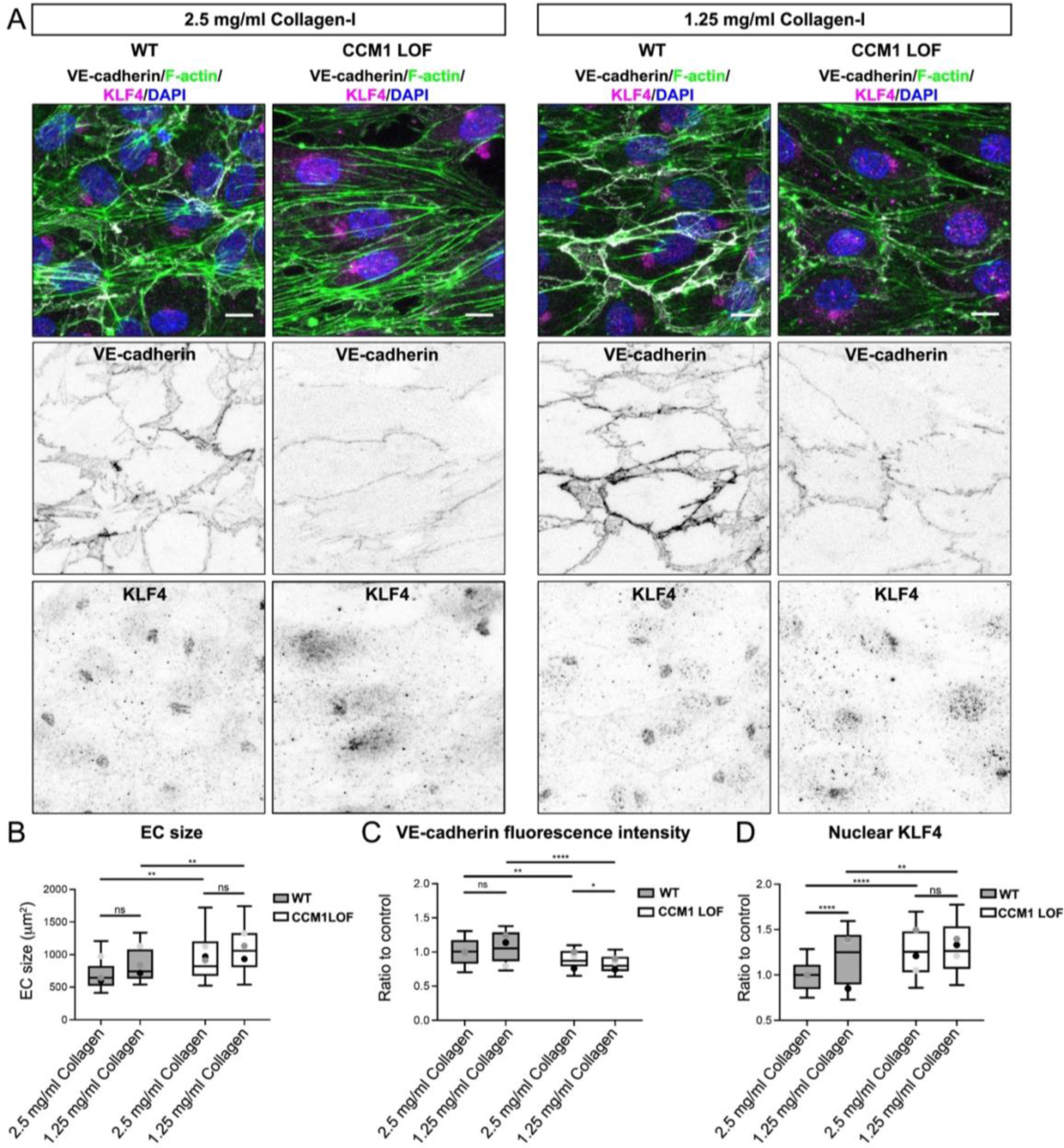
Decreasing collagen content and stiffness does not impact CCM severity. (A) Immunofluorescence of ECs in WT and CCM1 LOF micro-vessels grown for 48h in either 2.5 mg/ml Collagen-I (left) or 1.25 mg/ml Collagen-I (right), and stained for VE-cadherin, Phalloidin (F-actin), KLF4 and DAPI. Scale bar: 10µm. (B) Quantification of cell size of ECs in WT and CCM1 LOF micro-vessels grown in either 2.5 mg/ml or 1.25 mg/ml Collagen-I. Box and whisker plot with mean value of each replicate represented as a dot with matching colours between WT and CCM1 LOF ECs. n=3 replicates; 2.5 mg/ml Collagen-I n=58 WT and n=49 CCM1 LOF ECs, 1.25 mg/ml Collagen-I n=51 WT and n=54 CCM1 LOF ECs, Mann Whitney test **p<0.005, ns=no significant difference. (C) Quantification of the VE-cadherin fluorescence intensity in ECs of WT and CCM1 LOF micro-vessels grown in either 2.5 mg/ml or 1.25 mg/ml Collagen-I. Box and whisker plot with mean value of each replicate represented as a dot with matching colours between WT and CCM1 LOF ECs. n=3 replicates; 2.5 mg/ml Collagen-I n=58 WT and n=49 CCM1 LOF ECs, 1.25 mg/ml Collagen-I n=51 WT and n=54 CCM1 LOF ECs, Student’s t-test ***p<0.001, **p<0.005, *p<0.05, ns=no significant difference. (D) Quantification of the amount of nuclear KLF4, based on overlapping signal from KLF4 and DAPI. Box and whisker plot with mean value of each replicate represented as a dot with matching colours between WT and CCM1 LOF ECs. n=3 replicates; 2.5 mg/ml Collagen-I n=159 WT and n=164 CCM1 LOF ECs, 1.25 mg/ml Collagen-I n=165 WT and n=132 CCM1 LOF ECs, Mann Whitney test ****p<0.0001, **p<0.005, *p<0.05, ns=no significant difference.

### CCM1 loss alters transcriptional profiles of HA metabolism genes

The main ECM constituent of the brain is the glycosaminoglycan Hyaluronic acid (HA) which underpins unique CNS properties^27,39^. HA a particularly interesting component of the ECM as HA can elicit distinct biological responses, based on the length of the polysaccharide^29–31^. Disruption of HA homeostasis has therefore been associated with a wide range of pathologies, especially in the context of inflammation^40–43^. In the vasculature, distinct forms of HA have been identified to be required for developmental neo-angiogenesis^44^ and to control EC integrity^45^. HA production and turnover therefore needs to be tightly controlled. This occurs through the combined action of HA synthases (HAS) and Hyaluronidases. Notably, *HAS* genes and Hyaluronidases are amongst the most dysregulated genes in CCM1 deficient lesions from mice^32^. To determine the scope of CCM1 induced changes in HA metabolism genes, we performed quantitative mRNA expression analysis of all known HA synthases and Hyaluronidases in CCM1 LOF and WT ECs (Figure 4). CCM1 LOF ECs exhibited a marked increase of *HAS2* expression, whereas *HAS3* was significantly down regulated. HAS2 is associated with the synthesis of high-molecular-weight (HMW) HA (>500 kDa), whereas HAS3 produces low-molecular-weight (LMW) HA (<500 kDa)^30,46,47^. Furthermore, the Hyaluronidases *HYAL1* and *HYAL2* were significantly upregulated in CCM1 LOF ECs. Since HYAL1 and HYAL2 digest HMW HA into HA polymers of approximately 20 kDa^30^ this combined expression data implies that LMW HA production is decreased and that HMW HA is present, yet excessively degraded into HA polymers by HYAL1 and HYAL2. Interestingly, such HA polymers have been reported to have a pro-inflammatory effect via combined TLR4 and CD44 receptor activity^48^. Notably, TLR4 activation has been identified as a pre-requisite of CCM formation^49^. Changes in HA homeostasis in the CCM microenvironment therefore could alter signalling capabilities and impact the progression of this disease.

**Figure 4:**
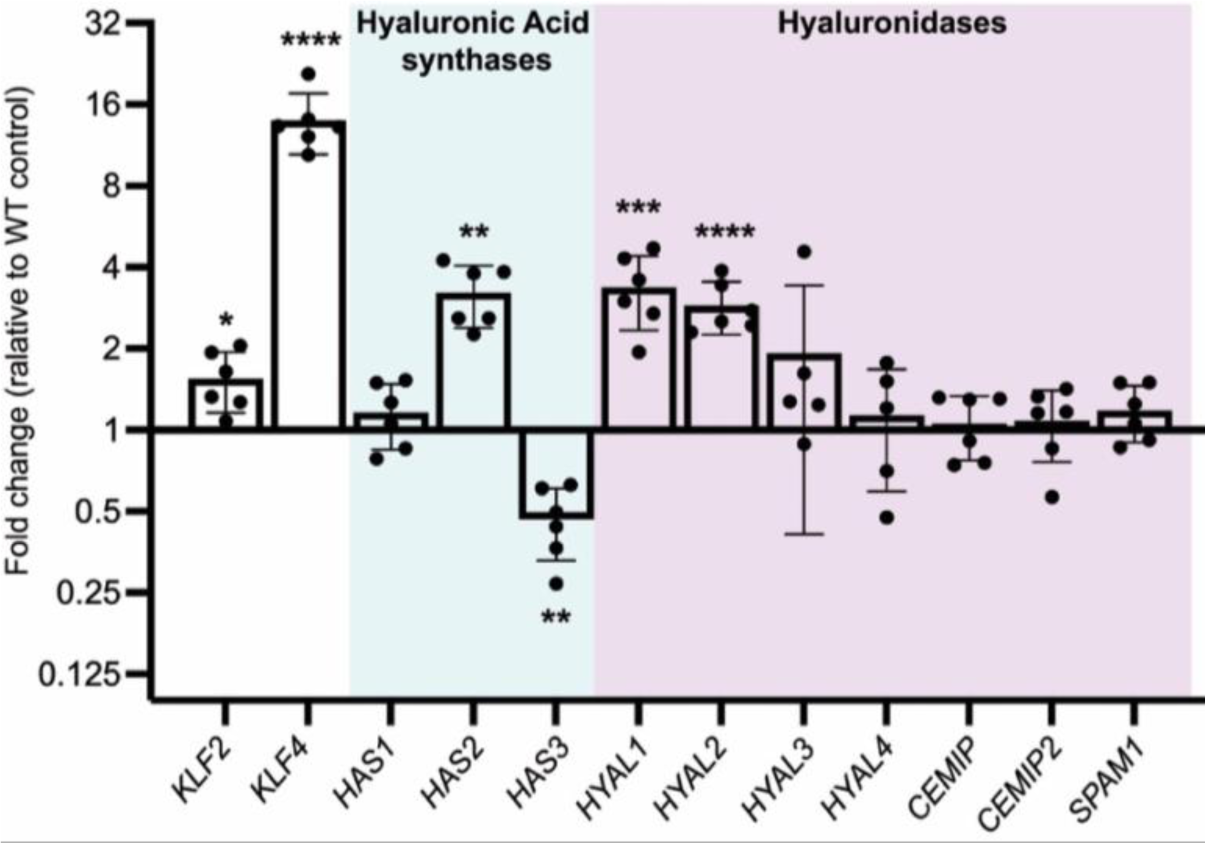
HA homeostasis in CCM1 LOF ECs. Quantitative RT-PCR analysis on mRNA isolated from WT and CCM1 LOF EC cells. Data presented as relative fold change to WT. Cyan background = Hyaluronic acid synthases and magenta background = Hyaluronidases. Expression of each gene was corrected relative to changes in the housekeeping gene *Hypoxanthine Phosphoribosyltransferase 1* (*HPRT*). Every dot-point represents a technical replicate (total of n=3 technical replicates) from n=2 independent biological replicates. Bars represent mean value. Error bars represent the standard deviation.

### HA of distinct molecular sizes can alter CCM cellular phenotypes

Based on the knowledge that HA can have strong effects EC biology, we used our 3D CCM model to test the phenotypic consequences when growing CCM vessels in matrices of predefined HA content. We first examined the impact of HA of distinct molecular weights, including short HA oligomers (<10kDa), LMW HA (41-65kDa), and HMW HA of 500-749kDa and 1-1.8MDa. We added 0.1% of each HA type to the standard Collagen-I ECM. The resulting ECMs were similar in terms of stiffness, collagen fibre architecture, and collagen pore size (Supplementary Figure 2D-F). CCM1 LOF and WT micro-vessels were grown in these distinct ECMs for 48 h and subsequently fixed for phenotypic analysis. By utilising VE-cadherin expression at the cell-cell junctions, we quantified EC size and identified that addition of LMW HA (41-65 kDa) significantly reduced the size of CCM1 LOF ECs, making them indistinguishable from WT control ECs (Figure 5A-B). This EC size rescue was specific to LMW HA, as CCM1 LOF ECs grown in the presence of HA oligomers or HMW HA were still significantly larger. However, HMW HA significantly restored VE-cadherin expression at junctions (Figure 5A, C). Notably, our expression analysis predicted that LMW and HMW HA might be reduced in the ECM due to changes HA synthesis and turnover when CCM1 is lost (Figure 4). Across all conditions nuclear KLF4 remained upregulated in CCM1 LOF ECs (Supplementary Figure 3A-B). Together, these results indicate that the combined action of LMW and HMW HA in the ECM might create a CCM protective microenvironment, whereby LMW and HMW HA likely govern EC size and junctional integrity through distinct pathways that act either downstream of KLF4 or in parallel. One of the main HA-receptors, CD44, is overexpressed on ECs in CCM^18,50,51^, which has been associated to a partial endothelial-to-mesenchymal (EndMT) of ECs in the lesions. In line with our observations, CD44 activation by HMW HA improves integrity of the EC barrier whilst shorter forms of HA exert the opposite effect when interacting CD44^45^. HA has also been reported to bind to Fibronectin^52^, thereby reducing outside-in integrin signalling capacity^53^. Since β1-Integrin overactivation has been associated with CCM, the inhibitory effect of LMW HA on EC enlargement might be a consequence of such a structural mechanism.

**Figure 5.**
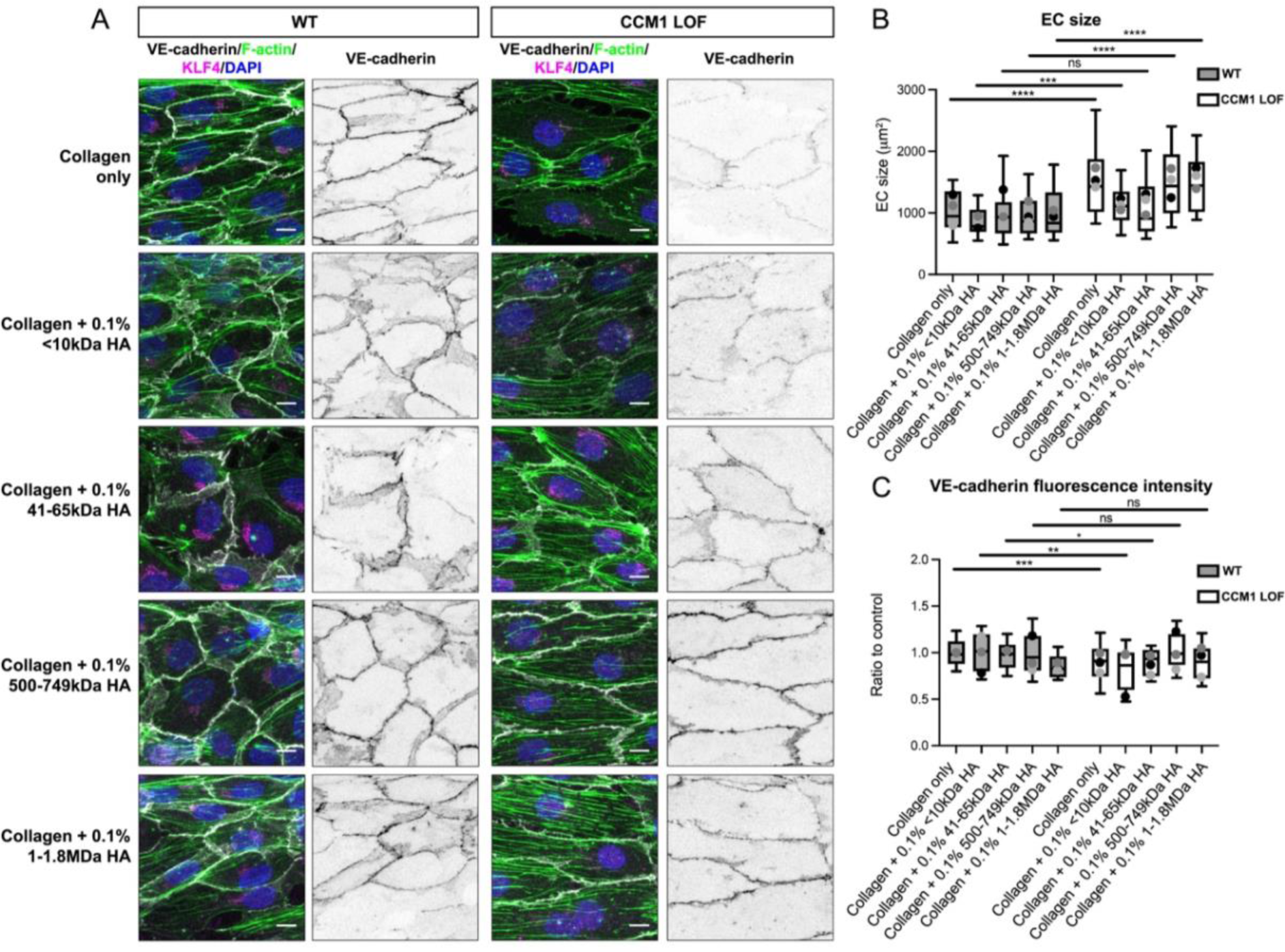
Impact of HA of defined molecular weight on CCM phenotype. (A) Immunofluorescence of WT and CCM1 LOF EC cells, seeded in 3D microfluidic devices in ECMs composed of either Collagen-I only, or Collagen-I with 0.1% of the Hyaluronic acid (HA) of incremental molecular weights. Tubes were grown for 48h post seeding and stained for VE-cadherin (white), Phalloidin (F-actin) (green), KLF4 (magenta) and DAPI (blue). Scale bar: 10µm. (B) Quantification of the cell size of WT and CCM1 LOF EC cells, measured based on VE-cadherin staining. Box and whisker plot with mean value of each replicate represented as a dot with matching colours between WT and CCM1 LOF ECs. n=3 replicates; 2.5 mg/ml Collagen-I n=66 WT and n=52 CCM1 LOF ECs, Collagen-I and 0.1% <10kD HA n=41 WT and n=36 CCM1 LOF ECs, Collagen-I and 0.1% 41-65kD HA n=45 WT and n=50 CCM1 LOF ECs, Collagen-I and 0.1% 500-749kD HA n=44 WT and n=42 CCM1 LOF ECs, Collagen-I and 0.1% 1-1.8 MDa HA n=44 WT and n=38 CCM1 LOF ECs. Mann Whitney test ****p<0.0001, ***p<0.001, ns=no significant difference. (C) Quantification of VE-cadherin expression at cell-cell junctions in WT and CCM1 LOF ECs. Box and whisker plot with mean value of each replicate represented as a dot with matching colours between WT and CCM1 LOF ECs. n=3 replicates; 2.5 mg/ml Collagen-I n=133 WT and n=107 CCM1 LOF ECs, Collagen-I and 0.1% <10kD HA n=41 WT and n=36 CCM1 LOF ECs, Collagen-I and 0.1% 41-65kD HA n=45 WT and n=50 CCM1 LOF ECs, Collagen-I and 0.1% 500-749kD HA n=43 WT and n=41 CCM1 LOF ECs, Collagen-I and 0.1% 1-1.8 MDa HA n=41 WT and n=42 CCM1 LOF ECs. Mann Whitney test ***p<0.001, **p<0.005, *p<0.05, ns = no significant difference.

To further test the effect of LMW HA (41-65 kDa) in controlling EC morphology, we increased the concentration of LMW HA in the ECM. At both concentrations of LMW HA we observed a significant decrease in EC size of the CCM1 LOF ECs (Figure 6A-B), further validating that LMW HA is sufficient to normalise the size of CCM1 LOF ECs and that EC size recovery is not concentration dependent. We next examined the functional requirement of LMW HA by degrading this HA in the ECM using a Hyaluronidase (HAse) from *Streptomyces hyalurolyticus*. This HAse acts on all types of HA, producing smaller HA degradation products of varied size^54^. Addition of HAse to hydrogels with 0.1% LMW HA, abolished EC size rescue, with CCM1 LOF ECs displaying increased size and reduced VE-cadherin, comparable to CCM1 LOF ECs grown in Collagen control ECMs (Figure 6A-C). This reversal was not apparent when adding HAse to 1% 41-65kDa HA ECMs (Figure 6A-C), suggesting that this amount of LMW HA could not be fully degraded by HAse and that sufficient LMW HA molecules remained to inhibit cellular expansion (Figure 6A-B). As we previously observed (Supplementary Figure 3), the effect of LMW HA on EC size was independent of KLF4 induction (Supplementary Figure 4 A-B).

**Figure 6.**
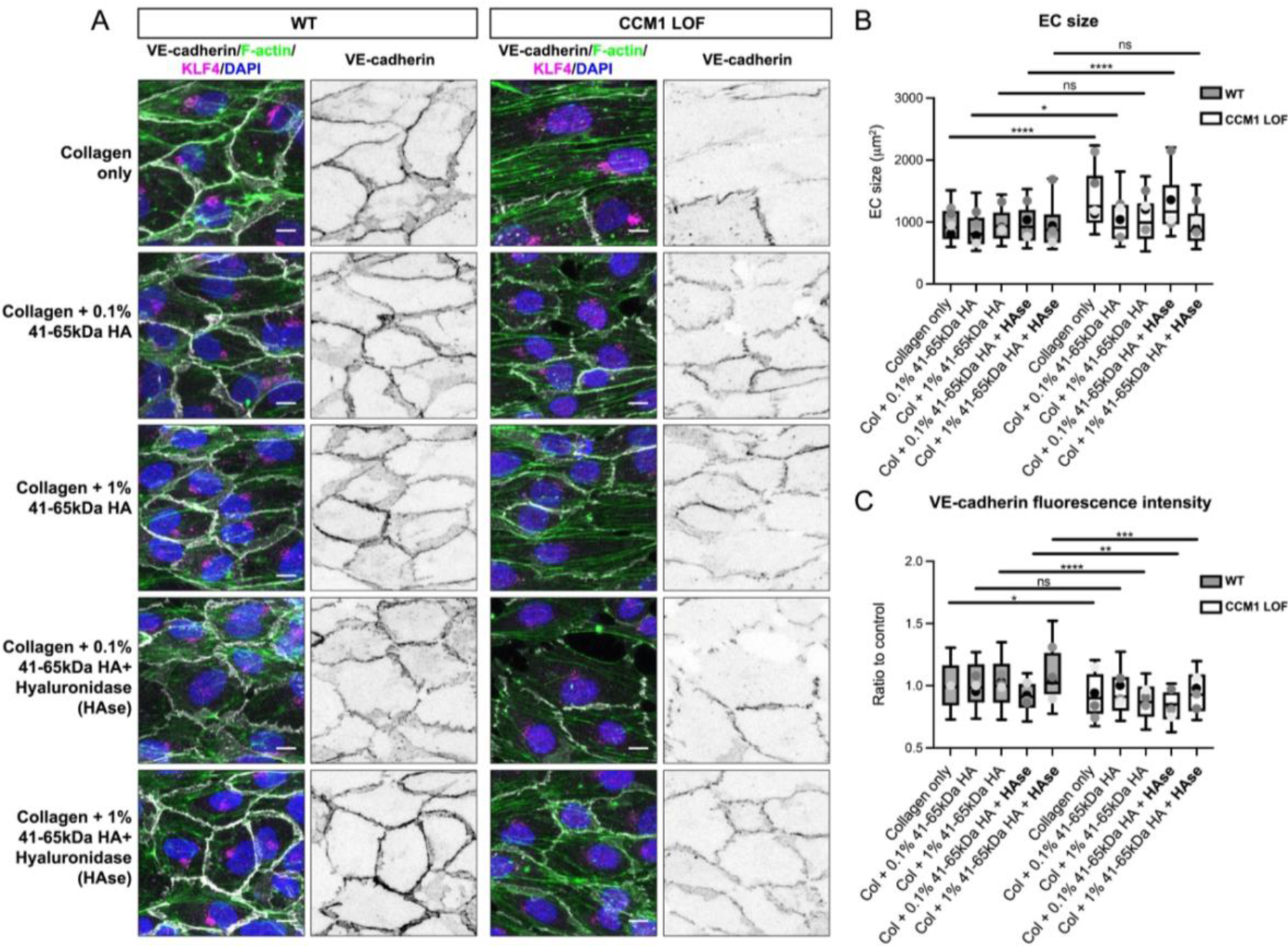
41-65kDa HA is required and sufficient to reduce CCM1 LOF EC size. (A) Immunofluorescence of WT and CCM1 LOF EC cells, seeded in 3D microfluidic devices. The composition of ECM is indicted for each condition. Cells were grown for 48h post seeding and stained for VE-cadherin (white), Phalloidin (F-actin) (green), KLF4 (magenta) and DAPI (blue). Scale bar: 10µm (B) Quantification of WT and CCM1 LOF EC cell size measured based on VE-cadherin staining. Box and whisker plot with mean value of each replicate represented as a dot with matching colours between WT and CCM1 LOF ECs. n=4 replicates; 2.5 mg/ml Collagen-I n=128 WT and n=118 CCM1 LOF ECs, Collagen-I and 0.1% 41-65kD HA n=140 WT and n=135 CCM1 LOF ECs, Collagen-I with 1% 41-65kD HA n=119 WT and n=125 CCM1 LOF ECs, Collagen-I with 0.1% 41-65kD HA and HAse n=72 WT and n=72 CCM1 LOF ECs, Collagen-I with 1% 41-65kD HA and HAse n=70 WT and n=86 CCM1 LOF ECs. Mann Whitney test ****p<0.0001, *p<0.05, ns=no significant difference. (C) Quantification of VE-cadherin expression at cell-cell junctions in WT and CCM1 LOF ECs. Box and whisker plot with mean value of each replicate represented as a dot with matching colours between WT and CCM1 LOF ECs. n=3 replicates; 2.5 mg/ml Collagen-I n=81 WT and n=75 CCM1 LOF ECs, Collagen-I and 0.1% 41-65kD HA n=94 WT and n=81 CCM1 LOF ECs, Collagen-I with 1% 41-65kD HA n=76 WT and n=72 CCM1 LOF ECs, Collagen-I with 0.1% 41-65kD HA and HAse n=72 WT and n=67 CCM1 LOF ECs, Collagen-I with 1% 41-65kD HA and HAse n=69 WT and n=85 CCM1 LOF ECs. Student’s t-test ****p<0.0001, ***p<0.001, **p<0.005, *p<0.05, ns=no significant difference.

Based on our observation that HMW forms of HA in the ECM improved EC integrity (Figure 5A, C), we next investigated whether a mixture of HMW and LMW HA fragments might be the most favourable to recover EC size and EC integrity. To test this, we generated ECMs with a combination of LMW HA and HMW HA fragments by adding HAse to HMW HA hydrogels. Without altering KLF4 expression (Supplementary Figure 4 C-D), both the size and VE-cadherin coverage of CCM1 LOF ECs was normalised in micro-vessels grown in the presence of LMW/HMW HA ECMs (Figure 7A-C), suggesting that it is the combined action of these HA forms that can prevent cellular phenotypes that are associated with CCM progression. Considering HA-based scaffolds are being developed as CNS therapeutics^55,56^, further research into the mechanism of distinct forms of HA in the CCM microenvironment might uncover new avenues to dampen CCM growth.

**Figure 7.**
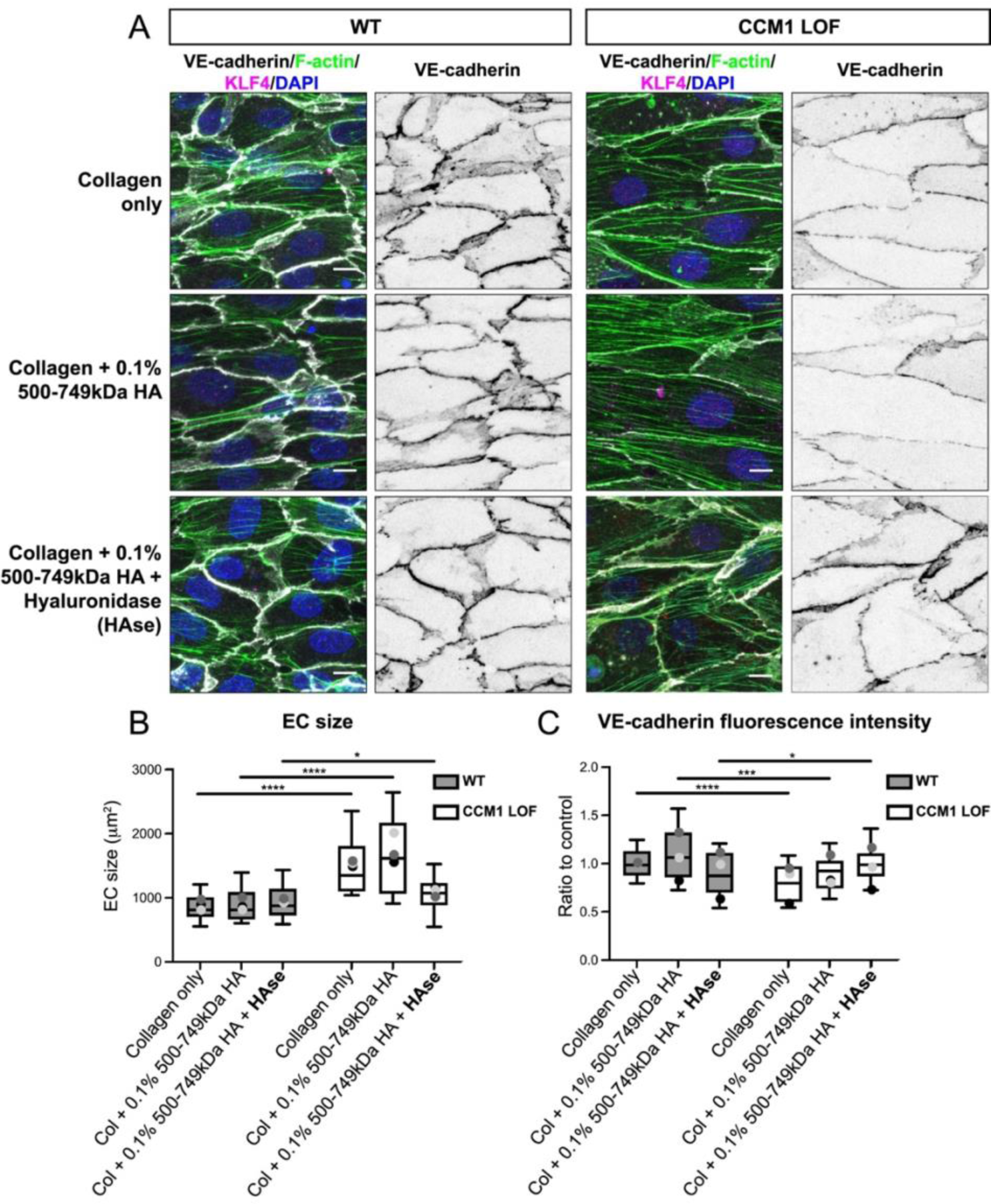
A combination of LMW and HMW HA in the ECM generates a CCM inhibitory environment. (A) Immunofluorescence of WT and CCM1 LOF ECs, seeded in 3D microfluidic devices. The composition of ECM is indicted for each condition. Cells were grown for 48h post seeding and stained for VE-cadherin (white), Phalloidin (Actin) (green), KLF4 (magenta) and DAPI (blue). Scale bar: 10µm. (B) Quantification of WT and CCM1 LOF EC size, measured based on VE-cadherin staining. Box and whisker plot with mean value of each replicate represented as a dot with matching colours between WT and CCM1 LOF ECs. n=3 replicates; 2.5 mg/ml Collagen-I n=46 WT and n=33 CCM1 LOF ECs, Collagen-I with 0.1% 500-749kD HA n=52 WT and n=38 CCM1 LOF ECs, Collagen-I with 0.1% 500-749kD HA and HAse n=38 WT and n=47 CCM1 LOF ECs. Mann Whitney test ****p<0.0001, *p<0.05, ns=no significant difference. (C) Quantification of VE-cadherin expression at cell-cell junctions in WT and CCM1 LOF ECs. Box and whisker plot with mean value of each replicate represented as a dot with matching colours between WT and CCM1 LOF ECs. n=3 replicates; 2.5 mg/ml Collagen-I n=44 WT and n=31 CCM1 LOF ECs, Collagen-I with 0.1% 500-749kD HA n=51 WT and n=36 CCM1 LOF ECs, Collagen-I with 0.1% 500-749kD HA and HAse n=38 WT and n=39 CCM1 LOF ECs. Student’s t-test ****p<0.0001, ***p<0.001, *p<0.05.

## CONCLUSION

In light of accumulating evidence that non-cell autonomous signals play a crucial role in CCM lesion progression^17,18,22,23^, we adapted a 3D micro-fluidic model^25,26^ to grow human CCM1 deficient micro-vessels. We validated that characteristic phenotypic changes such as EC enlargement, induction of KLF4 and reduced junctional integrity are miantained^13,15,57^. Reducing ECM stiffness did not worsen CCM phenotypes, suggesting that restrictiveness of CCMs is not solely explained by low tissue compliance. By transcriptional profiling, we identified that CCM LOF likely alters homeostasis of the main ECM component of the CNS, Hyaluronic acid (HA). By growing CCM1 LOF micro-vessels in ECMs enriched with distinctly sized HA, we uncovered that changes in HA composition of the ECM surrounding CCM lesions can impact cell phenotypes that would contribute to lesion development independent of the transcriptional inducer of CCMs, KLF4. In particular, supplementation with LMW and HMW HA in the ECM was most beneficial, with CCM1 LOF ECs being smaller with normalised VE-cadherin coverage. Since the CNS is an HA rich tissue, CCM induced changes to the HA equilibrium are likely to be exacerbated and might play a role in the CNS restrictiveness of this disease.

Overall, our findings highlight the effectiveness of 3D micro-vessel modelling in examining the role of ECM components in CCM. This approach will be applicable to research aimed to investigate similar non-cell autonomous mechanisms in other vascular malformations and diseases, such as lymphatic malformations and aneurysms.

## METHODS

### Reagents

Primary antibodies: mouse α-VE-cadherin (WB 1:1000; sc-9989; Santa Cruz), rabbit α-KRIT1 (CCM1) (WB 1:1000; ab196025; Abcam), rabbit α-GAPDH (WB 1:5000; 2118; Cell Signalling), mouse α-VE-cadherin, conjugated to Alexa 647 (IF 1:250; 561567; Becton Dickinson), rabbit α-KLF4 (IF 1:250; 4038; Cell Signalling), rabbit α-phosphorylated Paxillin (Y118) (IF 1:1000; 69363; Cell Signalling), mouse α-phosphorylated Myosin Light Chain (S19) (IF 1:250; 3675; Cell Signalling), rabbit α-phosphorylated Myosin Light Chain (T18/S19) (IF 1:500; 3674; Cell Signalling), rabbit α-beta Catenin (IF 1:1000; C2206; Sigma Aldrich), Phalloidin, conjugated to Alexa 488 (IF 1:500; A12379; Thermo Fischer), Phalloidin, conjugated to Alexa 647 (IF 1:500; ab176759; Abcam). For Western blotting, secondary antibodies, horseradish peroxidase (HRP)-conjugated goat α-mouse (Sigma Aldrich A4416; 1:10 000) and HRP-conjugated goat α-rabbit (Sigma Aldrich A0545; 1:10 000) were used. For immunofluorescence, Alexa Fluor 488 conjugated goat α-mouse (Thermo Fischer Scientific A-11001; 1:1000) and Alexa Fluor 568 conjugated goat α-rabbit (Thermo Fischer Scientific A-11036; 1:1000) were used. Additional reagents used: R&D Systems Culturex 3D Culture Matrix Rat Collagen I (In vitro technologies; RDS344702001), Poly-L-Lysine (Sigma Aldrich; P8920), Dulbecco’s Modified Eagle’s Medium (Sigma Aldrich; D5648), PDMS (AIBN; 04019862), Sodium Hyaluronate (<10kDa) (Lifecore Biomedical; HA5K-1), Sodium Hyaluronate (41-65kDa) (Lifecore Biomedical; HA40K-1), Sodium Hyaluronate (500-700kDa) (Lifecore Biomedical; HA700K-1), Sodium Hyaluronate (1-1.8MDa) (Lifecore Biomedical; HA15M-1), Hyaluronidase from *Streptomyces hyalurolyticus* (Sigma Aldrich; H1136).

### Cell Culture

Human umbilical vein endothelial cells (HUVECs) were purchased from LONZA (C2519A) and were cultured according to the supplier’s recommendations, using EBM-2 basal medium (CC-3156), supplemented with EGM-2 SingleQuots supplements (CC-4176). Cells were trypsinized using Trypsin-EDTA (LONZA; CC-5012) and maintained for up to five passages. Human Embryonic Kidney (HEK)-293T cells were cultured in DMEM (Thermo Fischer Scientific; 11995073), containing 10% fetal bovine serum (Thermo Fischer Scientific; 10099141) and 100 U/ml penicillin and streptomycin (Thermo Fischer Scientific; 15070063). Cells were trypsinized, using Trypsin-EDTA (Thermo Fischer Scientific; 15400054). All cell lines were cultured in a humidified 37°C incubator with 5% CO_2_.

### Lentivirus production

The guide (g)RNA sequence targeting *Homo sapiens KRIT1* (5’-GTATTCCCGAGAATTGAGACTGG-3’) was selected based on the Zhang lab online gRNA prediction tool (http://crispr.mit.edu/). The gRNA was subsequently cloned in lentiCRISPRv2 vector (Addgene; 52961), flanked by Esp3I (New England Biolabs; R0734) restriction sites. Lentiviral particles were produced in HEK 293T cells, which were co-transfected with the gRNA containing lentiCRISPRv2 vector together with the packaging vectors pMDLg/pRRE (Addgene; 12251), pRSV-Rev (Addgene; 12253) and pMD2.G (Addgene; 12259) using polyethyleneimine (Sigma Aldrich; 764604) as a transfection reagent. Supernatants containing viral particles were collected 48 and 72 h post transfection, concentrated with Lenti-X concentrator (Clontech; 631232) and used to infect HUVECs. Transduced HUVECs were selected with 1 ug/ml Puromycin (Sigma Aldrich; P8833).

### Generating 3D micro-vessels

Microfluidic devices were made as previously described^25,26^, using 2.5 mg/ml Collagen (In vitro technologies; RDS344702001) as a standard extracellular matrix. When indicated, HA polysaccharides were pre-mixed in the ECM at the specified concentrations. Hyaluronidase from *Streptomyces hyalurolyticus* (Sigma Aldrich; H1136) was used at 5 U/ml. HUVECs were close to confluency on plastic plated prior to seeded into the ECM and were maintained on a rocker in a humidified 37C° incubator with 5%CO_2_ to form tight tubes.

### Immunofluorescence and imaging

Cells, grown on glass cover slips were fixed in 4% formaldehyde solution in PBS for 10 min. Permeabilization was done, using 0.3% Triton X100 in PBS. Cells were blocked in a blocking buffer, consisting of 10% goat serum (Sigma Aldrich; G6767) in PBS for 1h at room temperature. Primary and secondary antibodies were diluted in blocking buffer in the above indicated dilutions and incubated for 1h each with 6 washing steps with PBS for 10 min in between. Final washing steps were carried out with the addition of 4′,6-diamidino-2-phenylindole. Cover slips were mounted on microscope slides using Mowiol. Fluorescent still images were acquired at a Zeiss AxioImage M2 upright microscope, equipped with a 63X N.A. 1.4 Oil objective and an Axiocam 506. Single plane confocal images were acquired a Zeiss Axiovert 200 Inverted microscope with an LSM 880 confocal scanner, equipped with a 63X N.A. 1.4 Oil objective. Collagen fibres were imaged on a Leica DMi8 inverted microscope with a SP8 Galvo and resonant confocal scanner, equipped with a 40X N.A. 1.1 Water objective. Microfluidic devices were fixed sequentially in 1% formaldehyde, containing 0.05% Triton for 90 sec, followed by 4% formaldehyde for 30 min at 37 C°. Tubes were permeabilised with 0.5% Triton for 10 min at 37C° and blocked for 4 h in 10% goat serum in PBS at 4 C° prior to staining. Z-stack image acquisition was performed on a confocal laser scanning microscope (Zeiss Axiovert 200 inverted microscope with LSM 710 Meta Confocal Scanner) using 40X NA 1.1 water immersion objective and 1 um Z step size.

### Western Blotting

Cell pellets were lysed in a lysis buffer containing 50mM Tris (pH 8.0), 150mM NaC, 1% Triton-X100, 10% Glycerol and protease inhibitors cocktail (Sigma #04693116001). Protein concentration was measured using the Bradford method^58^, prior to running the Western blots^59^. Blocking was done in a 3% BSA solution in PBS, containing 1mM EDTA and 0.05% Tween-20. Primary and secondary antibodies were diluted in the blocking solution and used according to the indicated concentrations with washing steps done in Tris-based saline with Tween 20 (TBST). Signals, emitted from the horseradish peroxidase were visualized by enhanced chemiluminescence (ECL) (BioRad #1705060) and imaged with Chemidoc (BioRad).

### Rheology

Rat tail collagen gels (In vitro technologies; RDS344702001) of different concentrations (1.25-2.5 mg/mL, pH 8-9, DMEM) and composites of collagen (2.5 mg/mL) and hyaluronan (1-10 mg/mL) of different molecular weights (<10 kDa - 1.8 MDa; LifeCore Biomedical) were analysed using an Anton Paar MCR-502 WESP rheometer. Samples were gelated *in situ* between plates by transferring the samples to a pre-cooled (5 °C) bottom stainless-steel plate (Anton Paar) using setting a gap of 0.5 mm with a stainless-steel parallel plate PP-25 (25 mm diameter, Anton Paar). Samples were trimmed and a thin layer of silica oil was placed around the sample. Gelation was induced with a temperature ramp from 5°C to 37°C (1 °C min^-1^) and further incubated for 2 hours at 37°C. Changes in the viscoelastic properties of the sample (G’ and G” (Pa)) were monitored by applying oscillatory shear at a constant strain of 0.5 % and frequency of 1 Hz.

### Quantitative RT-PCR

Cells were harvested directly in Buffer RLT with β-mercaptoethanol and RNA processed directly using the RNeasy Mini Kit (Qiagen), with on-column DNase digestion according to manufacturer’s instructions. RNA concentration was measured using a NanoDrop spectrophotometer and an equal starting concentration of RNA was for each sample was used for reverse transcription. Reverse transcription was performed using Superscript III Reverse Transcriptase (ThermoFisher) with Oligo dT priming. Quantitative PCR was performed using SYBR green reagent (Applied Biosystems) on a QuantStudio 7 Flex Real-Time PCR System (ThermoFisher) in 384 well plates (Applied Biosystems) and relative gene expression was determined using the change-in-threshold (2^-DDCT^) method, using Hypoxanthine Phosphoribosyltransferase 1 (HPRT) as a housekeeping control.

**Table I:**
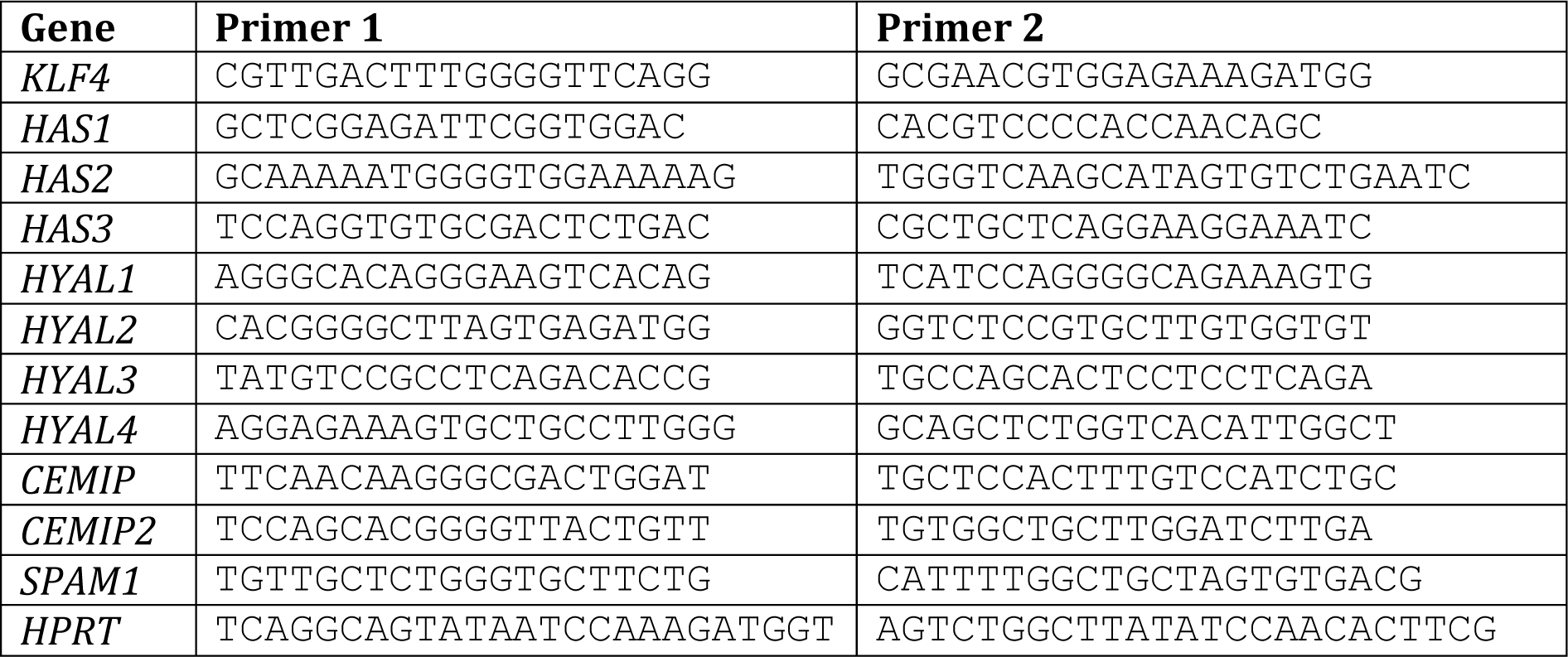
qPCR primers used:

### Statistical analysis

We performed all statistical analysis using Prism 9 (GraphPad). All quantifications were based on data acquired from at least three replicates. D’Agostino-Pearson test was applied to test normal distribution of the data points. When the data were normally distributed a Student’s t-test was used for comparison of two means. When the data did not follow a normal distribution, a Mann–Whitney test was used for comparison of two means. The threshold for significance was taken as p < 0.05.

## ACKNOWLEDGEMENTS

This work was performed in part at the Queensland node of the Australian National Fabrication Facility (ANFF), a company established under the National Collaborative Research Infrastructure Strategy to provide nano and microfabrication facilities for Australia’s researchers. Imaging was performed in the Australian Cancer Research Foundation’s Dynamic Imaging Facility at IMB (established with the generous support of the ACRF). We would also like to acknowledge Dr. Nicholas Condon for writing the Fiji scripts used for quantifications and Daniel Pelichowski for software support.

## ETHICS APPROVAL STATEMENT

Ethics approval not required

## AUTHOR CONTRIBUTIONS STATEMENT

Conceptualisation: T.E.Y., M.A.E.M., A.E.R., J.L., A.K.L.; Methodology: T.E.Y., M.A.E.M., T.E., J.B.T.; Validation: T.E.Y., M.A.E.M., A.K.L.; Formal analysis: T.E.Y., M.A.E.M., T.E., A.K.L.; Investigation: T.E.Y., M.A.E.M., T.E; Resources: J.B.T., L.I.L., S.J.S., A.E.R., B.M.H., C.S.C., J.L., A.K.L.; Data curation: T.E.Y., M.A.E.M., T.E., L.I.L., J.L., A.K.L.; Writing – original draft: T.E.Y., M.A.E.M., S.J.S., J.L., A.K.L.; Visualisation: T.E.Y., M.A.E.M., A.K.L.; Supervision: L.I.L., S.J.S., A.E.R., B.M.H., C.S.C., J.L., A.K.L.; Project administration: A.K.L.; Funding acquisition: L.I.L., S.J.S., A.E.R., B.M.H., C.S.C., A.K.L.

## DATA AVAILABILITY

The data that support the findings of this study are available from the corresponding author upon reasonable request.

