## Supplementary Material and Movie Captions for "Hyaluronic acid turnover controls the severity of Cerebral Cavernous Malformations in bioengineered human micro-vessels"

### SUPPLEMENTARY FIGURES

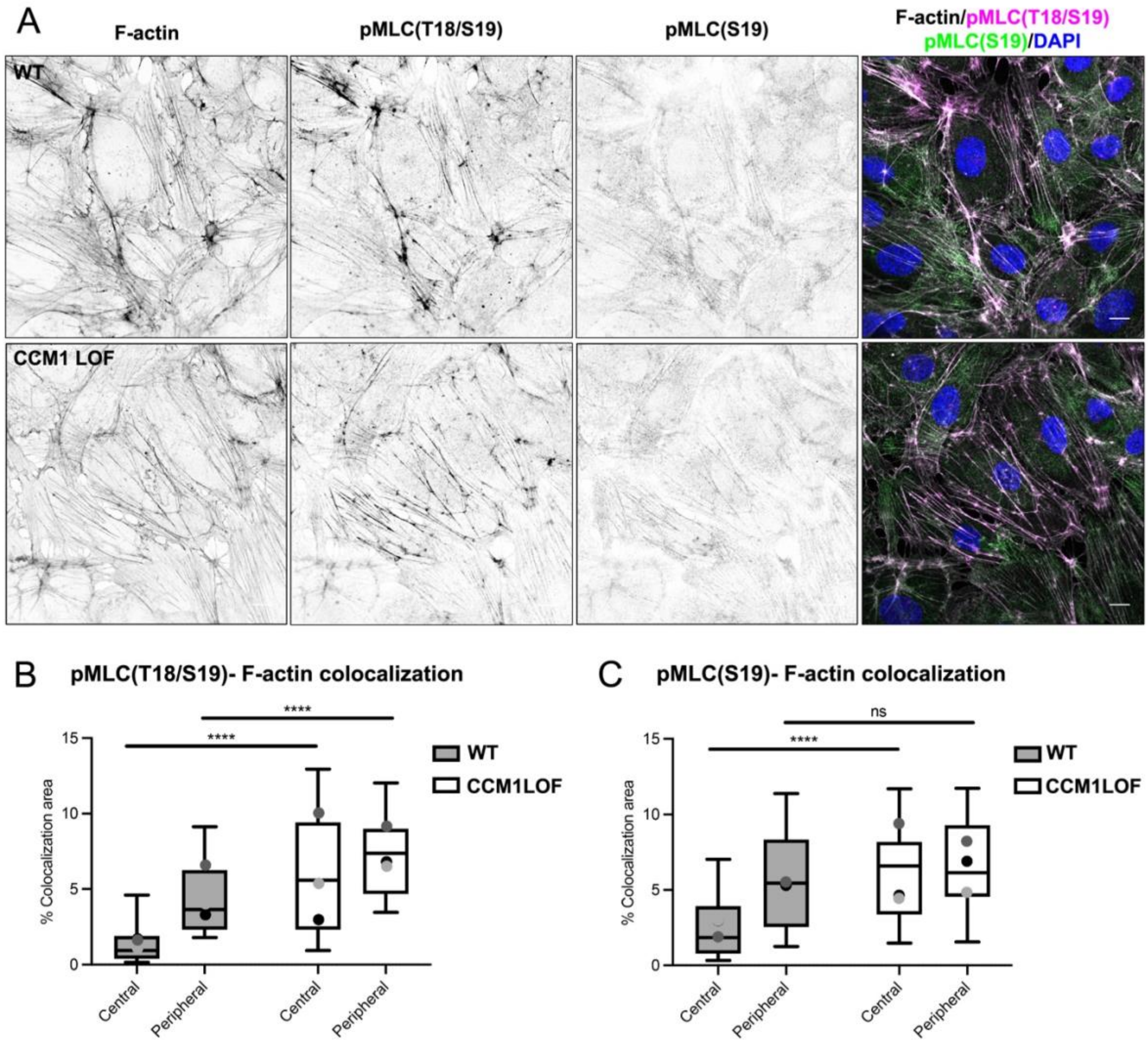


**Supplementary Figure 1: Induced pMLC expression and F-actin stress fibres in CCM1 LOF ECs**

(A) Immunofluorescence of WT and CCM1 LOF ECs, stained for F-actin (Phalloidin), phospho-MLC (T18/S19), phospho-MLC (S19) and DAPI. Scale bar: 10μm.

(B) Quantification of pMLC (T18/S19) and Phalloidin (F-actin) colocalization in either the central or the peripheral region of the ECs. Box and whisker plot with mean value of each replicate represented as a dot with matching colours between WT and CCM1 LOF ECs. n=3 replicates; n=40 WT and n=38 CCM1 LOF ECs. Mann Whitney test ****p<0.0001.

(C) Quantification of pMLC (S19) and Phalloidin (F-actin) colocalization in either the central or the peripheral region of the ECs. Box and whisker plot with mean value of each replicate represented as a dot with matching colours between WT and CCM1 LOF ECs. n=3 replicates; n=43 WT and n=38 CCM1 LOF ECs. Mann Whitney test ****p<0.0001, ns=no significant difference.

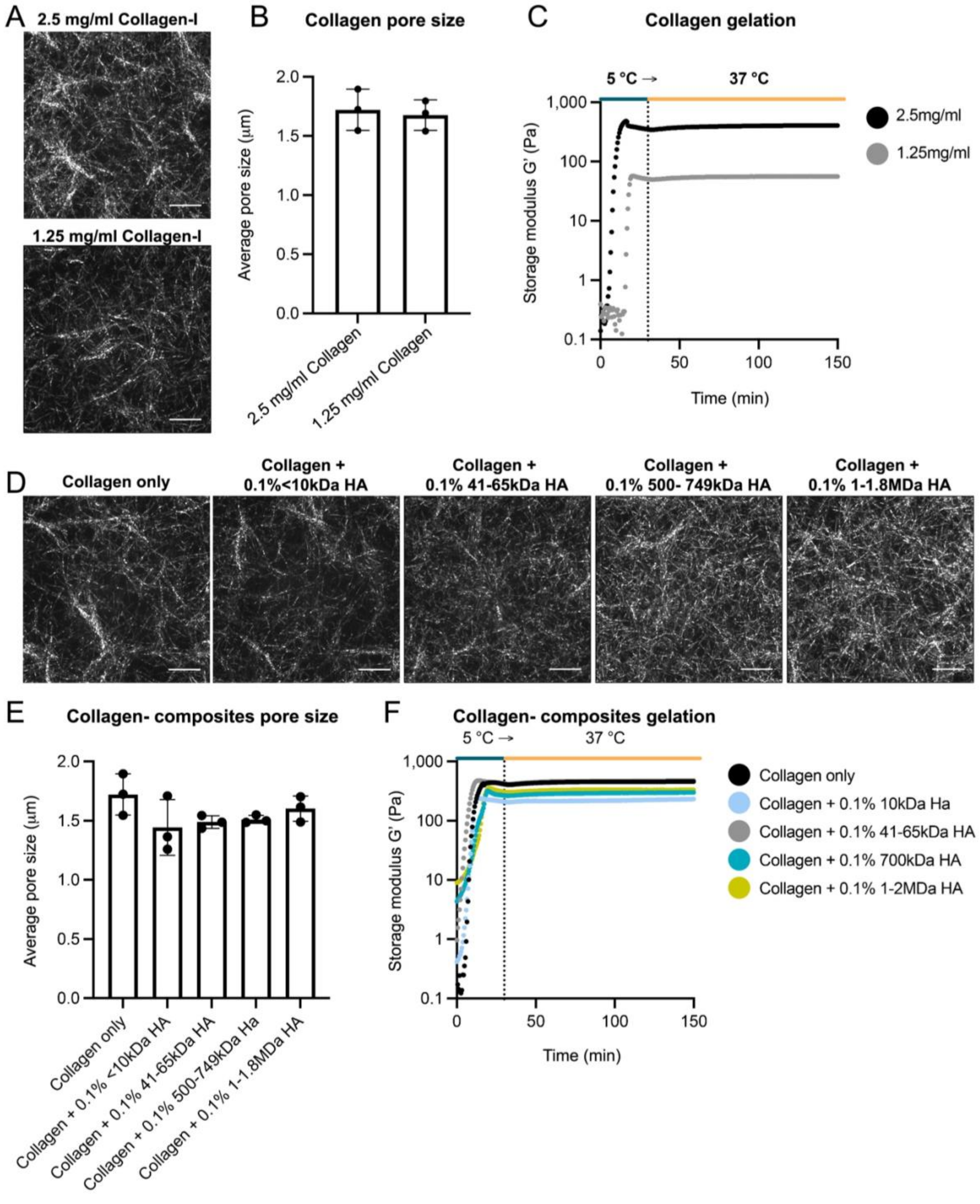


**Supplementary Figure 2: Mechanical and structural properties of Collagen hydrogels and Collagen-HA composites.**

(A) Maximum projections of confocal images of Collagen-I hydrogels, polymerised at 37°C, at 2.5 mg/ml (top) and 1.25 mg/ml (bottom). Scale bar: 10μm.

(B) Quantification of average pore size of 2.5 mg/ml and 1.25 mg/ml Collagen-I hydrogels, n=3 replicates.

(C) Rheological analysis of the storage modulus of the material ($G'$ (Pa)) for 2.5 mg/ml and 1.25 mg/ml Collagen-I hydrogels.

(D) Maximum projections of confocal images of Collagen-I hydrogels, polymerised at 37°C, with 2.5 mg/ml Collagen-I only (left) and Collagen-HA composites, containing 0.1% HA of defined molecular weights. Scale bar: 10μm.

(E) Quantification of average pore size of 2.5 mg/ml Collagen-I and Collagen-HA composites, n=3 replicates.

(F) Rheological analysis of the storage modulus of the material ($G'$ (Pa)) for 2.5 mg/ml Collagen-I only (left) and Collagen-HA composites, containing 0.1% HA of defined molecular weights.

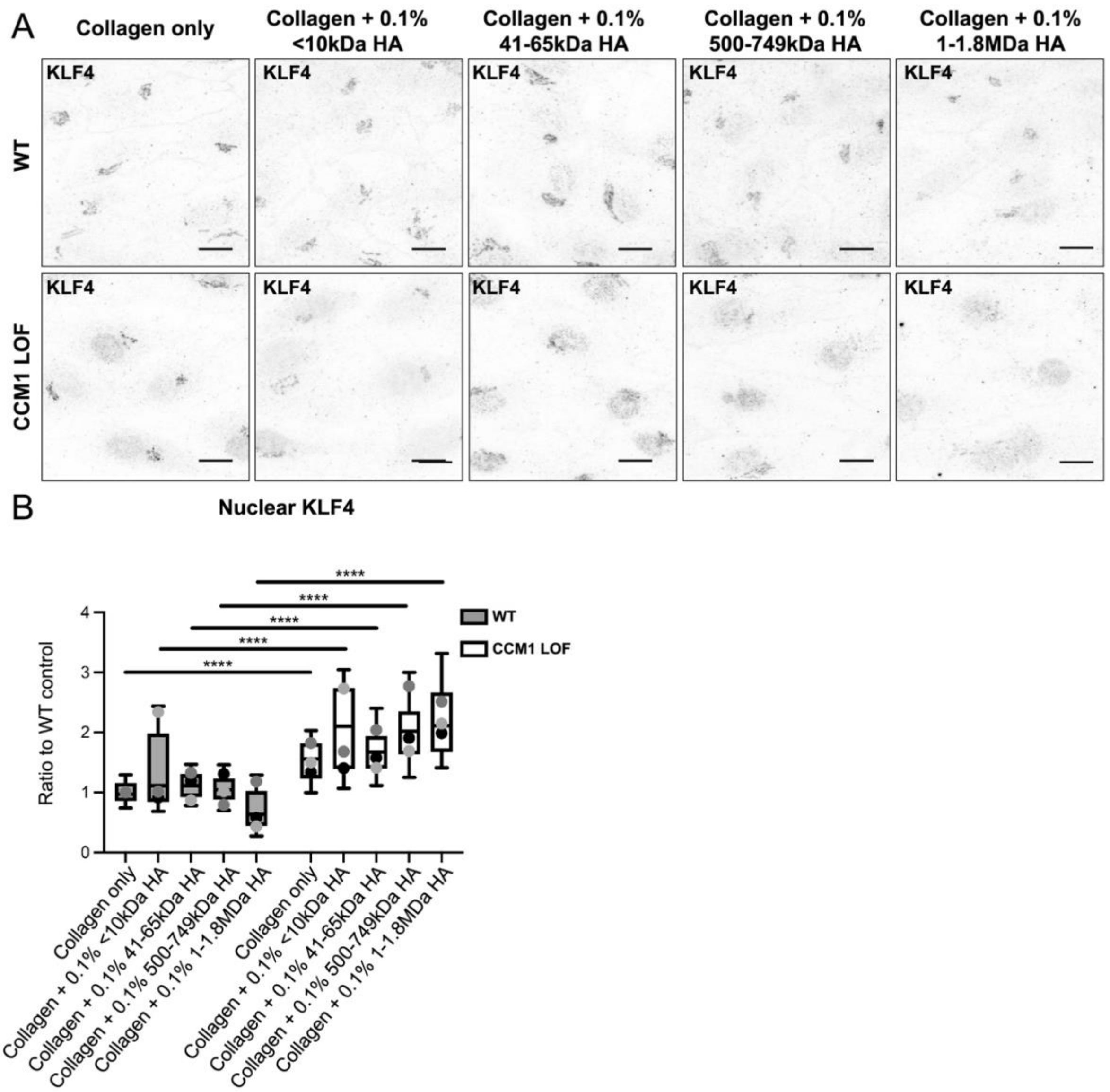


**Supplementary Figure 3: Distinct HA environments do not induce changes in ectopic KLF4 expression.**

(A) Supplementary to Figure 5. KLF4 expression only (grey). Scale bar: 10μm.

(B) Quantification of nuclear KLF4 expression, based on overlapping signal from KLF4 and DAPI. Box and whisker plot with mean value of each replicate represented as a dot with matching colours between WT and CCM1 LOF ECs. n=3 replicates; 2.5 mg/ml Collagen-I n=132 WT and n=108 CCM1 LOF ECs, Collagen-I and 0.1% <10kD HA n=153 WT and n=110 CCM1 LOF ECs, Collagen-I and 0.1% 41-65kD HA n=123 WT and n=118 CCM1 LOF ECs, Collagen-I and 0.1% 500-749kD HA n=157 WT and n=117 CCM1 LOF ECs, Collagen-I and 0.1% 1-1.8 MDa HA n=155 WT and n=141 CCM1 LOF ECs. Mann Whitney test ****$p<0.0001$.

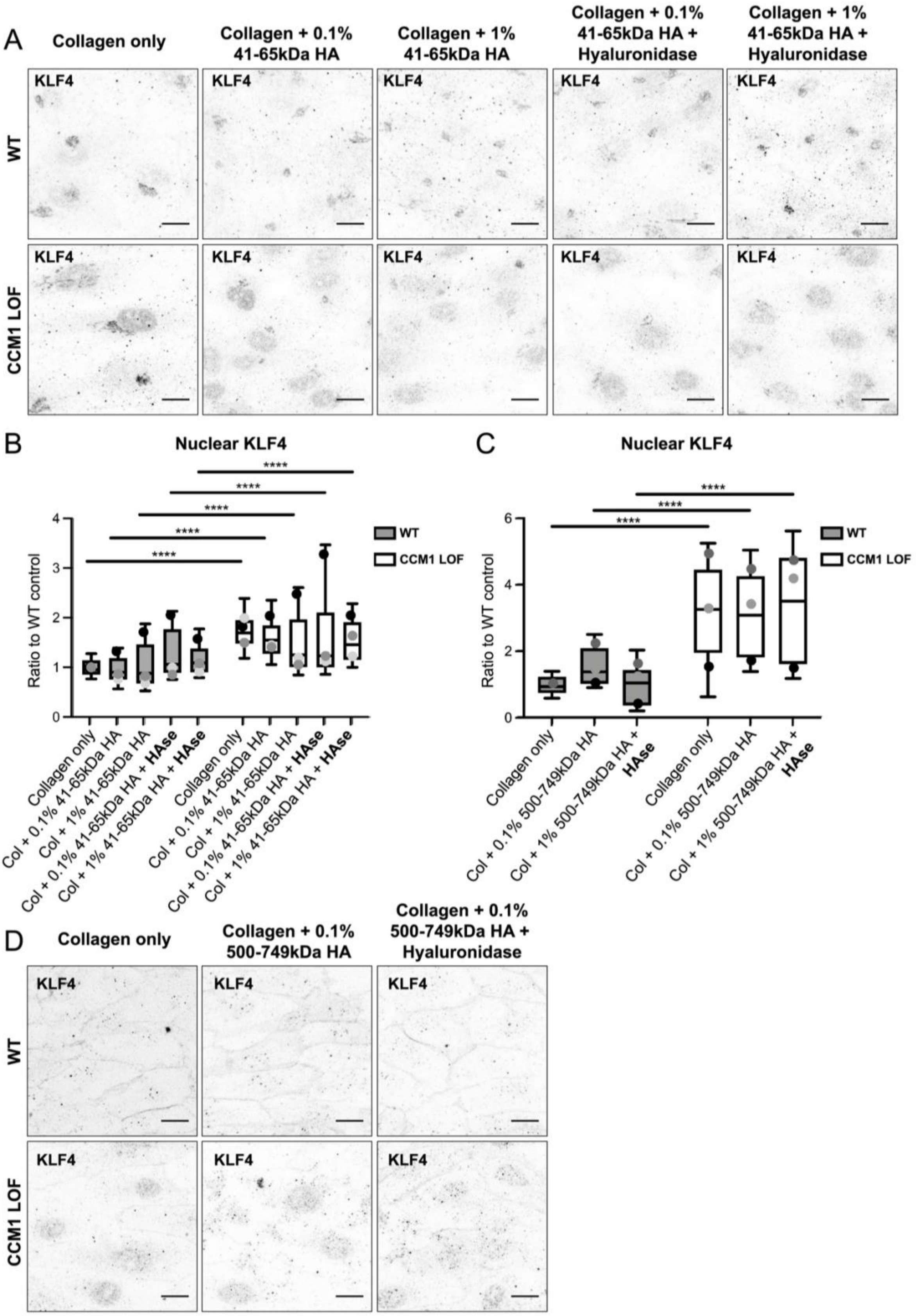
A
Collagen only
Collagen + 0.1% 41-65kDa HA
Collagen + 1% 41-65kDa HA
Collagen + 0.1% 41-65kDa HA + Hyaluronidase
Collagen + 1% 41-65kDa HA + Hyaluronidase
WT
CCM1 LOF
KLF4
B
Nuclear KLF4
Ratio to WT control
WT
CCM1 LOF
****
Collagen only
Col + 0.1% 41-65kDa HA
Col + 1% 41-65kDa HA
Col + 0.1% 41-65kDa HA + HAse
Col + 1% 41-65kDa HA + HAse
C
Nuclear KLF4
Ratio to WT control
Col + 0.1% 500-749kDa HA
Col + 1% 500-749kDa HA + HAse
D
Collagen only
Collagen + 0.1% 500-749kDa HA
Collagen + 0.1% 500-749kDa HA + Hyaluronidase

**Supplementary Figure 4: Increases in HA concentration or HA degradation do not induce changes in ectopic KLF4 expression.**

(A) Supplementary to Figure 6. KLF4 expression only (grey). Scale bar: 10 μm.

(B) Quantification of nuclear KLF4 expression, based on overlapping signal from KLF4 and DAPI. Box and whisker plot with mean value of each replicate represented as a dot with matching colours between WT and CCM1 LOF ECs. n=3 replicates; 2.5 mg/ml Collagen-I n=145 WT and n=101 CCM1 LOF ECs, Collagen-I and 0.1% 41-65kD HA n=145 WT and n=144 CCM1 LOF ECs, Collagen-I with 1% 41-65kD HA n=120 WT and n=149 CCM1 LOF ECs, Collagen-I with 0.1% 41-65kD HA and HAse n=139 WT and n=134 CCM1 LOF ECs, Collagen-I with 1% 41-65kD HA and HAse n=151 WT and n=156 CCM1 LOF ECs. Mann Whitney test ****p<0.0001, ***p<0.001.

(C) Quantification of nuclear KLF4 expression, based on overlapping signal from KLF4 and DAPI. Box and whisker plot with mean value of each replicate represented as a dot with matching colours between WT and CCM1 LOF ECs. n=3 replicates; 2.5 mg/ml Collagen-I n=168 WT and n=118 CCM1 LOF ECs, Collagen-I with 0.1% 500-749kD HA n=176 WT and n=109 CCM1 LOF ECs, Collagen-I with 0.1% 500-749kD HA and HAse n=173 WT and n=181 CCM1 LOF ECs. Mann Whitney test ****p<0.0001.

(D) Supplementary to Figure 7. KLF4 expression only (grey). Scale bar: 10 μm.

**Supplementary Movie 1: WT vessel in 3D microfluidic device.**

3D reconstruction of confocal Z-stacks of WT micro-vessels, grown in 2.5mg/ml Collagen-I based ECM. Scale bar: 10µm.

**Supplementary Movie 2: CCM1 LOF vessel in 3D microfluidic device.**

3D reconstruction of confocal Z-stacks of CCM1 LOF micro-vessels, grown in 2.5mg/ml Collagen-I based ECM. Scale bar: 10µm.
